# Engineering water exchange is a safe and effective method for magnetic resonance imaging in diverse cell types

**DOI:** 10.1101/2023.11.07.566095

**Authors:** Austin D.C. Miller, Soham P. Chowdhury, Hadley W. Hanson, Sarah K. Linderman, Hannah I. Ghasemi, Wyatt D. Miller, Meghan A. Morrissey, Chris D. Richardson, Brooke M. Gardner, Arnab Mukherjee

**Affiliations:** Biomolecular Science and Engineering Graduate Program, University of California, Santa Barbara, CA 93106, USA; Department of Molecular, Cellular, and Developmental Biology, University of California, Santa Barbara, CA 93106, USA; Department of Chemical Engineering, University of California, Santa Barbara, CA 93106, USA; Department of Bioengineering, University of California, Santa Barbara, CA 93106, USA; Department of Chemistry, University of California, Santa Barbara, CA 93106, USA; Neuroscience Research Institute, University of California, Santa Barbara, CA 93106, USA

**Author notes:** Correspondence should be addressed to AM.

**Keywords:** aquaporin-1, diffusion, MRI, reporter gene, unfolded protein response, cell physiology

## Abstract

Aquaporin-1 (Aqp1), a water channel, has garnered significant interest for cell-based medicine and in vivo synthetic biology due to its ability to be genetically encoded to produce magnetic resonance signals by increasing the rate of water diffusion in cells. However, concerns regarding the effects of Aqp1 overexpression and increased membrane diffusivity on cell physiology have limited its widespread use as a deep-tissue reporter. In this study, we present evidence that Aqp1 generates strong diffusion-based magnetic resonance signals without adversely affecting cell viability or morphology in diverse cell lines derived from mice and humans. Our findings indicate that Aqp1 overexpression does not induce ER stress, which is frequently associated with heterologous expression of membrane proteins. Furthermore, we observed that Aqp1 expression had no detrimental effects on native biological activities, such as phagocytosis, immune response, insulin secretion, and tumor cell migration in the analyzed cell lines. These findings should serve to alleviate any lingering safety concerns regarding the utilization of Aqp1 as a genetic reporter and should foster its broader application as a noninvasive reporter for in vivo studies.

## Introduction

Genetic reporters are indispensable tools for synthetic biology and basic research because they allow molecular events to be quantitatively tracked in living cells by measuring physical signals, such as light. However, light penetration is hindered by photon scattering and absorption, limiting the effectiveness of optical reporters for noninvasive deep-tissue imaging in vertebrates^1^. The need to overcome this limitation has motivated efforts to engineer genetic reporters for tissue-penetrant imaging modalities including ultrasound^2–5^, nuclear imaging^6–8^, and MRI^9–25^. Among these modalities, MRI is unsurpassed for generating high-resolution images across large volumes of tissue at any depth, without radiation hazards. Conventional genetic reporters for MRI take advantage of the magnetic properties of metals to create a contrast between engineered cells and the surrounding tissue^16–18,20,21,26–29^. However, the permeability of metals through the cell membrane and anatomical barriers, such as the blood-brain barrier, is non-uniform, which makes it challenging to accurately measure reporter activity in vivo. Furthermore, the exposure of cells to metals increases the risk of toxicity^30–34^. We recently introduced a metal-free, fully autonomous (i.e., no external agents required), single-gene MRI reporter based on human aquaporin-1 (Aqp1), a channel protein that facilitates the rapid and selective exchange of water molecules across the plasma membrane^35^. Water diffuses faster in tissues consisting of cells engineered to express Aqp1 than in tissues comprising wild-type cells (**Fig. 1a**). This increase in water diffusivity can be quantitatively imaged using an MRI technique known as diffusion-weighted imaging^36^.

**Figure 1:**
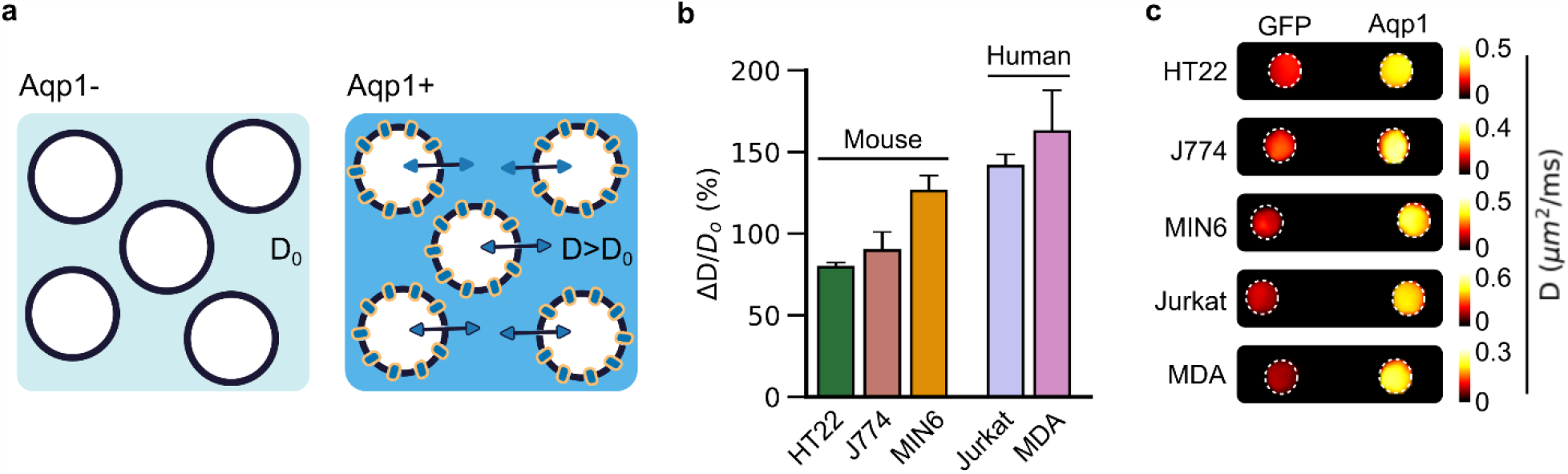
Diffusion-weighted imaging of Aqp1-expressing cells. a, Illustration of the Aqp1 contrast mechanism. Heterologous expression of Aqp1 increases the baseline water diffusivity (D_o) of cells by facilitating rapid water exchange across the cell membrane. The larger diffusivity of Aqp1-expressing cells (D) makes them observable using diffusion-weighted MRI. b, Percent increase in diffusivity (ΔD/D_o) of Aqp1-expressing cells relative to control cells transduced to express GFP. Error bars represent the s.e.m. from n = 4-8 independent biological replicates. c, Diffusion maps of axial cross-sections of pellets of Aqp1- and GFP-expressing cells. Each voxel in the diffusion map represents absolute diffusivity, estimated from a voxel-wise regression of the first-order decay in signal intensity with diffusion-weighting (i.e., effective b-value). The diffusion map is denoised using a median filter and displayed using a linear 8-bit color map whose lower and upper limits denote diffusivity in µm2/ms. All MRI data were acquired at 7 T, using a diffusion time of 300 ms.

Recent studies have expanded Aqp1 beyond our initial proof-of-concept by demonstrating that Aqp1 can be used to track tumor-specific gene expression^37^, trace neural connectivity^38^, and generate brain-wide maps of astrocyte populations in mice^39^. However, the effects of Aqp1 on cell health and function remain largely unknown. For example, it is unclear whether driving an increase in water diffusion imposes particular risks to cell health or whether introducing Aqp1 through heterologous expression induces metabolic burden. Additionally, it is important to assess the risk of incorrect folding of overexpressed membrane proteins in the endoplasmic reticulum (ER) and its consequences such as ER stress and cell damage. Therefore, a deeper understanding of Aqp1-related toxicity and perturbations to cell function is needed to ensure the safety and efficacy of Aqp1-based imaging techniques and make informed decisions regarding their use in deep-tissue imaging.

In this study, we evaluated the potential risks associated with using Aqp1 as a molecular reporter. We tested Aqp1 for its ability to sufficiently enhance water diffusivity in several different cell lines of neuronal, macrophage, pancreatic, T-lymphocyte, and tumor origin from both mouse and human lineages, confirming its broad utility as an MRI reporter. We probed the potential for adverse effects on cell physiology using biophysical and biochemical assays to measure changes in cell mass, volume, shape, viability, and caspase activity in Aqp1-expressing cells relative to GFP-controls. We also probed Aqp1-expressing cells for ER stress by measuring relative changes in genetic markers of the unfolded protein response (UPR). Finally, we tested Aqp1-expressing cells for their ability to perform cell type-specific functions, such as immune response, phagocytosis, insulin secretion, and mig ration.

## Results

### Aqp1 expression enhances water diffusivity in various cell types

Our first objective was to determine whether Aqp1 expression increased diffusivity in diverse mammalian cell types. To this end, we used lentiviral infection to express Aqp1 from a strong constitutively active promoter (EF1α) in five cell lines: HT22 (hippocampal), J774A.1 (macrophage), Jurkat (T lymphocyte), MDA-MB-231 (triple-negative breast cancer), and MIN6 (insulinoma). We selected these cell lines to represent a variety of biological and biophysical distinctions (**Table S1**), including different cell lineages, adherent and suspension cell types, male and female, human and mouse origins, and a range of baseline diffusivities (**Supplementary Fig. 1a**). We co-expressed Aqp1 with a fluorescent reporter (GFP) using an internal ribosome entry site (IRES) to generate uniform populations of Aqp1-expressing cells by fluorescence-activated cell sorting (FACS) (**Supplementary Fig. 2**). We used diffusion-weighted MRI to measure the increase in water diffusivity in Aqp1-expressing cells relative to cells lacking Aqp1, but similarly transduced to express GFP. We observed a substantial increase in diffusivity in all cells, ranging from 79.4 ± 2.8 % in HT22 to 162.5 ± 25.2 % in MDA-MB-231 (**Fig. 1b, Supplementary Fig. 1b-f**). The lower fold-change in HT22 compared to other cell types arises from its faster baseline diffusion rate (**Supplementary Fig. 1a**). Taken together, these results suggest that Aqp1 expression is a viable method for imaging a broad range of mammalian cell types using diffusion-weighted MRI (**Fig. 1c**).

### Aqp1 expression does not alter cell mass, volume, sphericity, and viability

To assess whether Aqp1 expression and the ensuing increase in water diffusivity is a safe approach for imaging cells, we compared Aqp1-expressing cells to those expressing GFP in terms of dry mass, volume, and sphericity (viz. roundedness), key morphological descriptors that are often altered by adverse changes in cell physiology^40–46^. To measure these biophysical parameters, we used quantitative phase microscopy (**Fig. 2a**), a tomographic technique that extracts morphometric features by mapping the refractive index distribution within cells^47,48^. We observed no significant variations in dry mass (**Fig. 2b**), volume (**Fig. 2c**), and sphericity (**Fig. 2d**) between Aqp1- and GFP-expressing cells, indicating that Aqp1 expression does not interfere with key physiological mechanisms, including biosynthetic and degradative pathways, ionic homeostasis, and osmotic gradients that collectively regulate cell size and shape^49–52^. We also evaluated cell viability using standard assays based on MTT reduction and ATP content^53,54^, finding no major differences in Aqp1-expressing cells relative to GFP-transduced controls (**Fig. 2e,f**). Additionally, Aqp1 expression did not cause activation of executioner caspase-3/7 in any cell line, indicating no induction of programmed cell death (**Fig. 2g**). Taken together, these results indicate that overexpression of Aqp1 does not incur a metabolic or safety burden notably greater than that of GFP expression.

**Figure 2:**
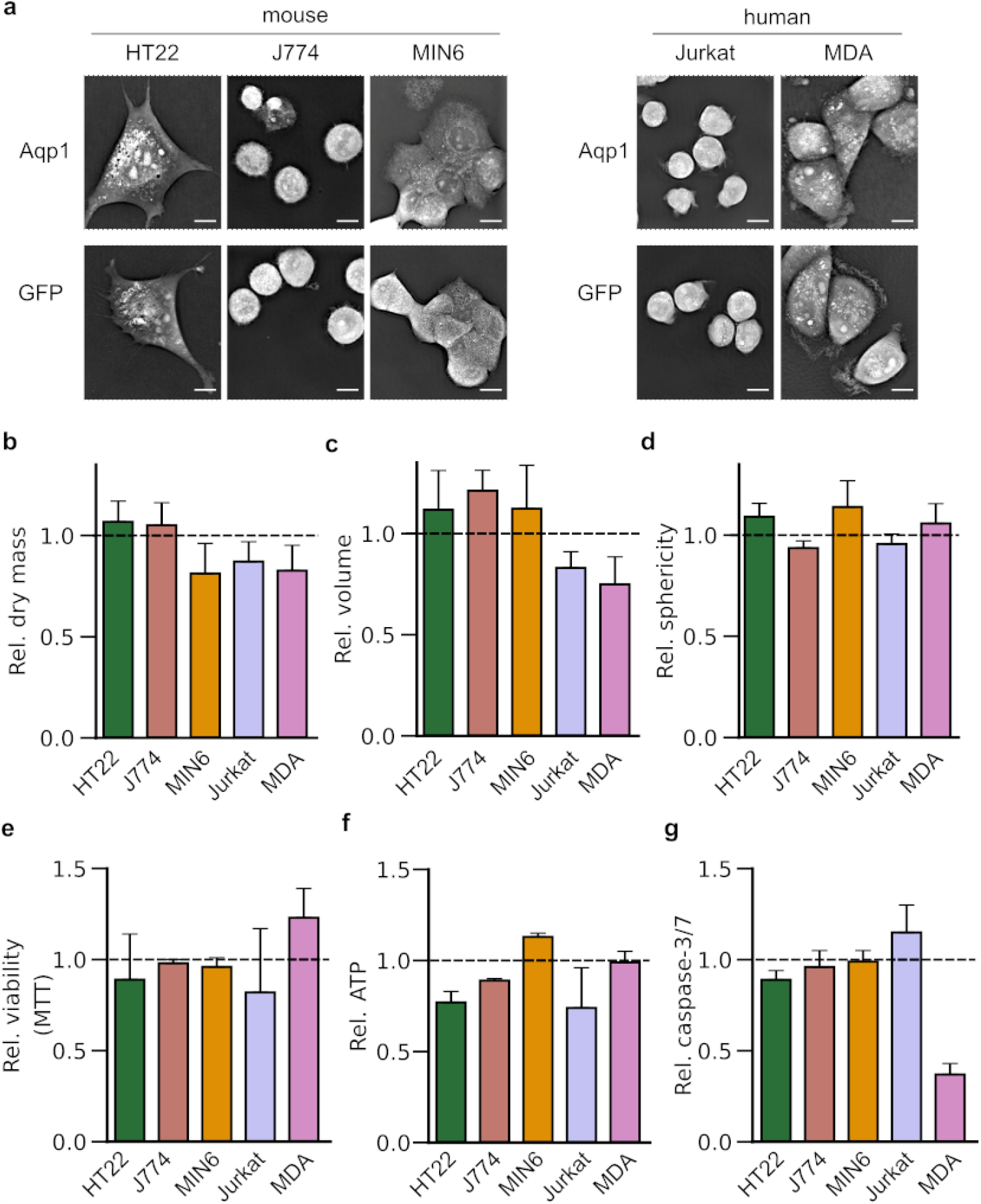
Effects of Aqp1 expression on cell morphology and viability. a, Representative quantitative phase microscopy images of cells engineered to express Aqp1 or GFP. Scale bar is 5 µm. Fold-change in b, dry mass, c, volume, and d, sphericity of Aqp1-expressing cells relative to GFP-cells measured using quantitative phase imaging. e, Viability of Aqp1-expressing cells relative to GFP-controls measured using the MTT assay. f, ATP levels in Aqp1-expressing cells relative to GFP. g, Caspase-3/7 activation in Aqp1-cells relative to GFP. The dashed lines denote the unit fold change representing no difference between GFP and Aqp1. Error bars in the phase imaging experiments represent the s.e.m. from n ≥ 3 images comprising 10-50 single cells. Error bars in the viability and caspase assays represent the s.e.m. from n ≥ 3 biological replicates.

### Aqp1 expression does not trigger the unfolded protein response

Cells activate the unfolded protein response (UPR) as an adaptive mechanism when the protein folding machinery of the ER is overwhelmed by the buildup of misfolded or unfolded proteins in the organelle^55,56^. Notably, overexpression of secreted and membrane-bound proteins, including antibodies^57^, GPCRs^58^, transporters^59^, and microbial opsins^60^, have been shown to trigger the UPR in cells. Accordingly, we sought to determine whether the UPR is induced by heterologous expression of Aqp1, a channel protein that must fold and mature inside the ER prior to reaching the cell membrane. We used qRT-PCR, to measure the fold-change (relative to GFP controls) in the levels of key UPR-associated genes^61^, including BiP, CHOP, and XBP1s. Additionally, we assessed changes in the expression of activating transcription factor 4 (ATF4), an ER chaperone (Grp94), and an E3 ubiquitin ligase (SYVN1) involved in ER-associated degradation of misfolded proteins^62^. We did not detect a significant increase in any UPR-associated gene in Aqp1-expressing cells (**Fig. 3a, Supplementary Fig. 3)**, with the exception of Grp94, which showed an approximately two-fold increase in expression in MIN6 cells (**Supplementary Fig. 3**). Next, we investigated whether Aqp1-expressing cells retained the ability to activate the UPR when exposed to ER stress by treatment with tunicamycin, a protein glycosylation inhibitor that leads to extensive accumulation of misfolded proteins in the ER^63^. Aqp1-expressing cells showed tunicamycin-dependent increase (relative to vehicle-treated cells) in levels of BiP, XBP1s, and CHOP (**Fig. 3b**), suggesting that Aqp1 expression does not impair the innate capacity of cells to respond to ER stress.

**Figure 3:**
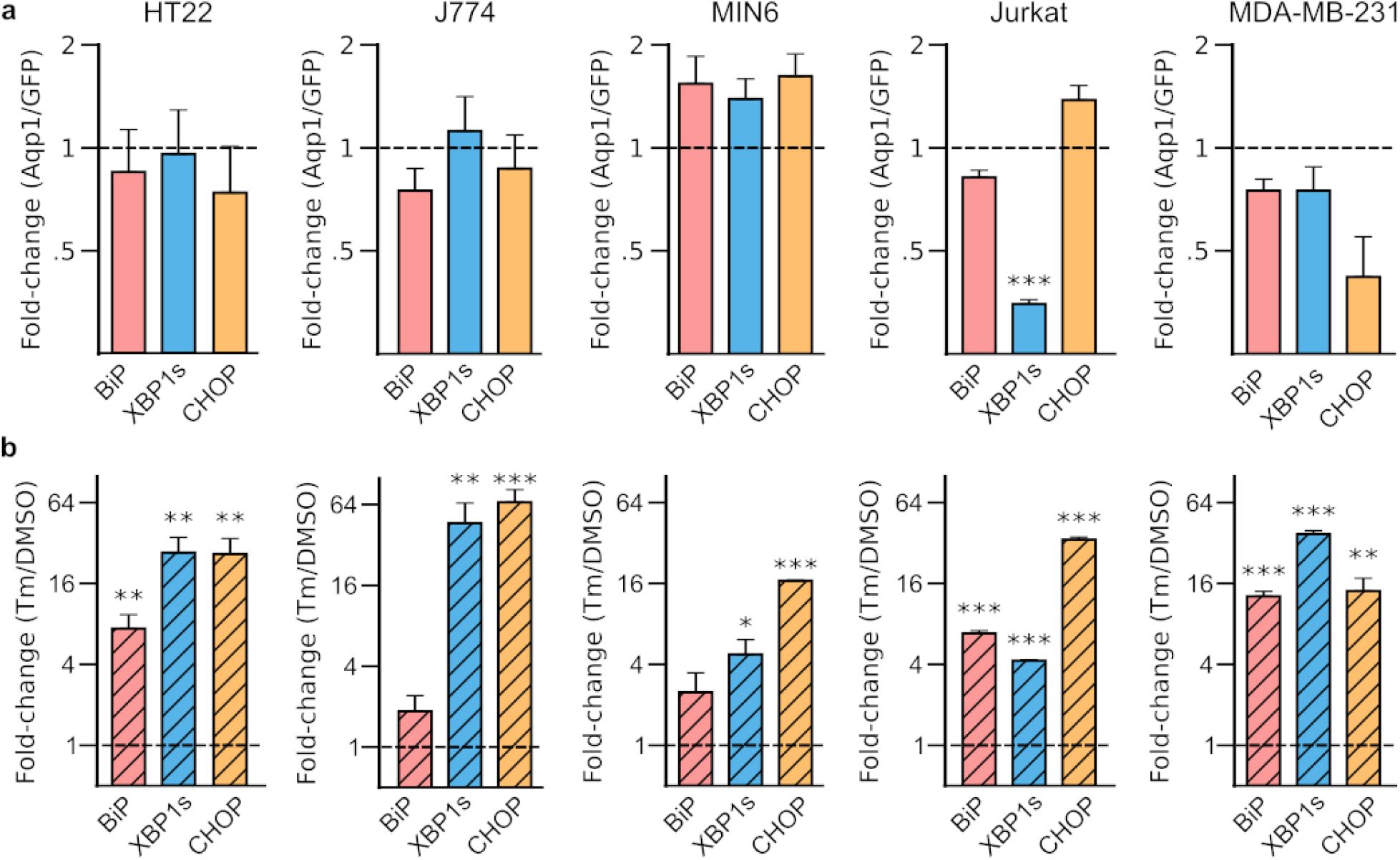
Effect of Aqp1 expression on the unfolded protein response. **a**, Fold changes in key UPR-associated genes, BiP, XBP1s, and CHOP in Aqp1-expressing cells relative to GFP controls. **b**, Fold changes in the BiP, XBP1s, and CHOP in Aqp1-expressing cells treated with 2.5 µg/mL tunicamycin for 4 h relative to vehicle-treated cells. Error bars represent the s.e.m. from *n* = 3 biological replicates. * *P*-value < 0.05, ** *P*-value < 0.01, *** *P*-value < 0.001. GAPDH and actin were used as housekeeping genes for the mouse and human cell lines, respectively.

### Aqp1 expression does not adversely affect cell type-specific functions

Having established that Aqp1 expression does not substantially perturb key descriptors of cell morphology, cytotoxicity, and ER stress, we wondered whether Aqp1 expression could interfere with functions in various cell types. To address this question, we used a panel of assays to test Aqp1-expressing cells for in vitro activities that model the in vivo function specific to a particular cell-type, including phagocytosis (J774A.1), insulin secretion (MIN6), CD25 expression (Jurkat), and metastatic invasion (MDA-MB-231). We excluded HT22 cells (neuronal) from this analysis, as a recent study showed that overexpression of Aqp1 in murine neurons caused no difference in the electrical properties of Aqp1-neurons compared with GFP-transduced neurons^38^. To assess the functionality of J774A.1 cells, we used silica beads coated with a supported-lipid bilayer to emulate the surface of cells. We incorporated biotinylated lipids in the membrane and incubated the beads with monoclonal anti-biotin IgG, a well-characterized “eat-me” signal^64^. We mixed the beads with Aqp1- and GFP-expressing cells and measure internalized beads using confocal microscopy as described in our earlier work^65^. Both Aqp1- and GFP-expressing cells showed similar phagocytic activities, which increased by two-fold following exposure to beads containing IgG in the lipid bilayer relative to cells incubated with IgG-free beads (**Fig. 4a,b**,**Supplementary Fig. 4**). These findings indicate that Aqp1 expression does not impede phagocytic activity in macrophages. Next, we assessed MIN6 functionality by measuring glucose-stimulated insulin secretion. In this assay, cells are stimulated to release insulin by elevating external glucose concentration, which is used to model pancreatic beta-cell function^66,67^. Both Aqp1- and GFP-expressing cells showed significant glucose-stimulated insulin secretion, with GFP-expressing cells having a modestly higher fold-change (2.2 ± 0.2 fold, mean ± s.e.m) than Aqp1-cells (1.6 ± 0.2 fold) (**Fig. 4c**). To test Jurkat cell function, we stimulated the T-cell receptors with anti-CD3/CD28 microbeads and used flow cytometry to measure increase in surface expression of CD25, a well-defined marker of T-cell activation^68,69^. As expected, both Aqp1- and GFP-expressing Jurkat cells showed a significant increase in CD25 surface expression (**Fig. 4d-e, Supplementary Fig. 5**); however, an approximately two-fold larger fraction of Aqp1-expressing cells upregulated CD25 compared to GFP-expressing cells (**Fig. 4f**). Finally, we studied the migration ability of MDA-MB-231 cells using a Matrigel-based assay^70^, which is commonly employed to assess tumor cell invasiveness in in vitro drug screening platforms. Specifically, we seeded cells in a Matrigel drop and used confocal imaging to measure the area occupied by the cells that migrated beyond the drop perimeter (**Fig. 4g**). Both Aqp1- and GFP-expressing cells exhibited migration profiles similar to those previously reported for wild-type MDA-MB-231 (**Fig. 4h**). Taken together, our findings suggest that Aqp1 expression does not significantly impede physiological responses, except for MIN6, where Aqp1 expression is associated with a statistically significant decrease in insulin release.

**Figure 4:**
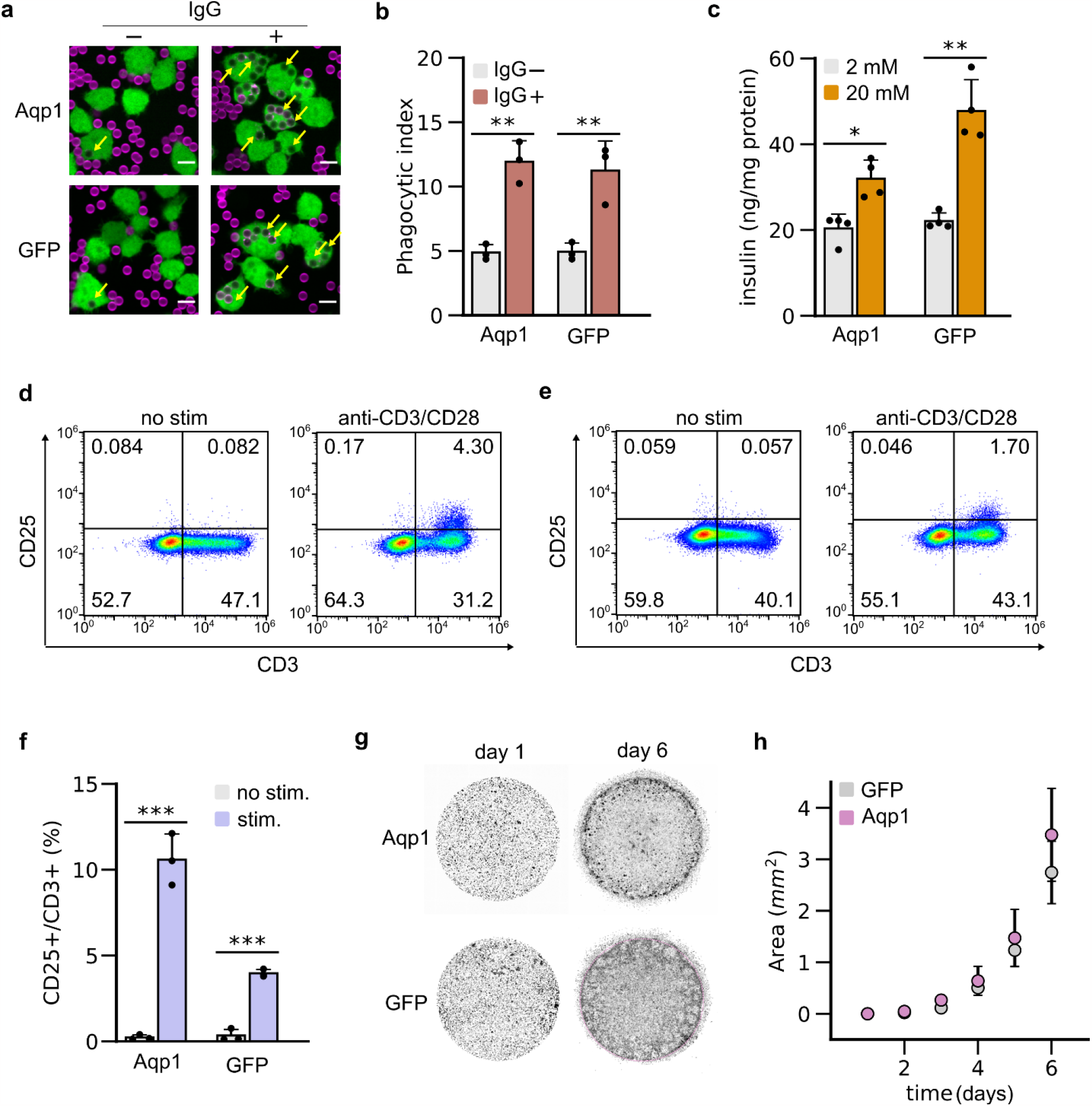
Effect of Aqp1 expression on specific cell functions. **a**, Representative images showing the engulfment of silica beads coated with a supported lipid bilayer (atto390, magenta) by Aqp1-IRES-GFP and GFP-expressing J774A.1 cells (GFP, green). Images were collected using a spinning disk confocal microscope with a 40X 0.95 NA Plan Apo air objective. Yellow arrows denote beads engulfed by the cells. Scale bar is 5 µm. **b**, Phagocytic activity of Aqp1- and GFP-cells following exposure to lipid-coated silica beads with or without IgG1κ incorporated in the supported lipid bilayer, for 45 min. Phagocytic index was determined by measuring the average att390 fluorescence per macrophage.**c**, Glucose-stimulated insulin secretion in Aqp1- and GFP-expressing MIN6 cells induced by exposing cells to 20 mM external glucose for 1 h. **d**, Flow cytometry analysis of CD25 and CD3 surface expression in Aqp1- and **e**, GFP-expressing Jurkat cells, either untreated or incubated with anti-human CD3/CD28 beads at a 4:1 bead-to-cell ratio for 24 h. The numbers in each quadrant denote the fraction of cells showing surface expression of CD3 (pan T-cell marker), CD25 (activation marker), both CD3 and CD25, or neither marker. **f**, Percentage of CD25^+^ cells relative to the total CD3^+^ population. **g**, Representative images showing the migration of Aqp1- and GFP-expressing MDA-MB-231 cells beyond the Matrigel perimeter on day 6. Wide-field images covering both the Matrigel drop and the surrounding media were acquired using a scanning confocal microscope with a 10 × 10 grid of 1024 × 1024 pixel tiles at 10X magnification. **h**, Increase in the total area occupied by invading MDA-MB-231 cells that migrate out of the Matrigel. Error bars represent the s.e.m. from *n* ≥ 3 biological replicates. * *P*-value < 0.05, ** *P*-value < 0.01, *** *P*-value < 0.001 (2-sided, t-test).

## Conclusions

With its ability to genetically encode diffusivity signals that can detected using standard MRI instrumentation and pulse sequences, Aqp1 holds promise for revealing deeper insights into biological processes in their native context. Accordingly, we were motivated to systematically study Aqp1’s effects on cell physiology and function to address concerns regarding how Aqp1 expression and increased membrane diffusivity potentially affect cell physiology. We showed that Aqp1 expression generates a substantial increase in diffusivity in various mouse and human cell lines, suggesting that the Aqp1 reporter generalizes to diverse cellular chassis. We found no adverse effects of Aqp1 expression on several biophysical and biochemical parameters, including dry mass, volume, shape, viability, apoptosis, and ER health. We further showed that Aqp1 expression did not hinder the specific functions of different cells, such as phagocytosis, immune activation, and tumor cell migration. Altogether, our findings should reassure safety concerns regarding Aqp1, encouraging its broad application as a molecular tool for deep-tissue imaging, similar to the widespread utilization of GFP and its derivatives for visualizing cultured cells and transparent organisms.

The current study has some limitations that should be taken into account. First, we evaluated the performance of Aqp1 in vitro using cultured cells, which provides a convenient platform to survey a range of biological and biophysical variables (**Table S1**), and are routinely employed for toxicology studies. In the future, validating Aqp1 safety through in vivo studies in small and large animal models would be an additional step forward. Second, our safety and toxicity tests were conducted under standard non-stressed cell culture conditions, and therefore may not accurately reflect the effects of Aqp1 in more stressful environments. In the future, Aqp1 could be “pressure tested” in environments where the cell experiences additional metabolic or biosynthetic burden, such as during hypoxia or when co-expressed with other membrane-bound genetic tools, for example channelrhodopsins or chimeric antigen receptors. Third, the study was conducted with cells that were cultured and passaged for 2-3 weeks after transduction. Therefore, the effects of prolonged (e.g. several weeks) Aqp1 expression on cell behavior are not fully unraveled. Further research should examine the effects of longer durations of Aqp1 expression to assess its suitability for longitudinal imaging paradigms. Lastly, we observed that Aqp1-expressing MIN6 cells exhibited elevated expression of the UPR marker, Grp94 (**Supplementary Fig. 3**), suggesting the potential presence of secretory burden associated with Aqp1 overexpression. These findings imply that cells with a high secretory burden, as is the case with insulin secretion, may be more susceptible to the effects of Aqp1 overexpression. However, further investigation is needed to gain a deeper understanding of the biological significance of these results. If necessary, the Aqp1-related secretory burden can be reduced by employing a weaker promoter or implementing a conditional expression system that restricts Aqp1 expression to a specified time window. Future research can explore these options to determine the optimal window for balancing Aqp1 expression, signal-to-noise ratio, and metabolic burden in cell types where sustained overexpression of Aqp1 appears to interfere with cell physiology.

In summary, our study contributes to the ongoing development of tissue-penetrant reporters by addressing potential safety concerns related to the use of Aqp1 as a reporter gene. We expect that our findings will have significant practical implications in several areas of basic and applied research, including synthetic biology, genetic medicine, immuno-oncology, and systems neuroscience, where the ability to track cells and transcriptional activity in intact living systems is a highly sought-after capability.

## Methods Reagents

Molecular biology reagents, including Q5® High-Fidelity 2X Master Mix, 1 kb Plus DNA ladder (100 bp-10 kb), and agarose gel electrophoresis loading dye were purchased from New England Biolabs (Ipswich, MA, USA). iScript™ cDNA synthesis reagents were purchased from Bio-Rad (Hercules, CA, USA). PowerUp™ SYBR™ Green Master Mix for quantitative RT-PCR was purchased from Thermo Fisher Scientific (Waltham, MA, USA). Agarose and SYBR™ Safe DNA stain were respectively purchased from GoldBio (St. Louis, MO, USA) and APExBIO (Houston, TX, USA). Reagents for plasmid extraction (mini- and midi-prep) and DNA purification were purchased from Promega (Madison, WI, USA) and New England Biolabs. Reagents for RNA extraction were purchased from Qiagen (Hilden, Germany). Chemically competent *E. coli* was obtained from New England Biolabs. Oligonucleotide primers for PCR amplification were designed using NEBuilder and ordered from Integrated DNA Technologies (Carlsbad, CA, USA).

HT-22 cell lines were purchased from Sigma-Aldrich (St. Louis, MO, USA). MIN6 cells were purchased from Thermo Fisher Scientific. All other cell lines were obtained from the American Type Culture Collection (ATCC). Dulbecco’s Modified Eagle Media (DMEM), Roswell Park Memorial Institute media (RPMI 1640), sodium pyruvate, and casein were purchased from Sigma-Aldrich (St. Louis, MO, USA). GlutaMAX™ (100X), penicillin-streptomycin (10^4^ units/mL penicillin, 10 mg/mL streptomycin), sterile phosphate buffered saline (PBS), sodium butyrate, CellTracker™ Green dye, TrypLE, and anti-human CD3/CD28 magnetic Dynabeads™ were purchased from Thermo Fisher Scientific. Fetal bovine serum (FBS) was purchased from R&D Systems (Minneapolis, MN, USA) and Thermo Fisher Scientific. Linear polyethyleneimine (25kDa) transfection reagent was purchased from Polysciences Inc. (Warrington, PA, USA). Lenti-X™ viral concentrator was purchased from Takara Bio (San Jose, CA, USA). RIPA cell lysis buffer and polybrene were purchased from Santa Cruz Biotechnology (Dallas, TX, USA).

Reagents for performing cytotoxicity and viability assays, namely CellTiter-Glo®, CellTiter 96® AQ_ueous_, Caspase-Glo® 3/7, and CellTiter-Blue® were purchased from Promega (Madison, WI, USA). Lipids were purchased from Avanti (Alabaster, AL, USA) and ATTO-TEC (Siegen, Germany). Silica beads were purchased from Bangs labs (Fishers, IN, USA). Monoclonal mouse anti-biotin IgG1κ was purchased from Jackson ImmunoResearch Labs (West Grove, PA, USA). All other monoclonal antibodies were purchased from Thermo Fisher Scientific. Matrigel® was purchased from Corning (NY, USA). Pierce™ BCA Protein Assay reagent were purchased from Thermo Fisher Scientific. Ultra-Sensitive Mouse Insulin ELISA kit was purchased from Crystal Chem (Elk Grove Village, IL).

## Molecular biology

The construction of the Aqp1-expressing lentiviral plasmid (pJY22) has been described in our previous study. An enhanced GFP reporter was co-expressed with Aqp1 using an internal ribosome entry site (IRES) to allow selection of stably transduced cells by fluorescence-activated cell sorting (FACS). To generate a control plasmid (pADM04), the GFP sequence was amplified using Q5® High-Fidelity 2X Master Mix and cloned by Gibson assembly in the same lentiviral vector backbone used for Aqp1 expression. All constructs were verified using Sanger DNA sequencing (Genewiz, San Diego, CA, USA).

### Cell culture and engineering

Cells were routinely cultured at 37 °C in a humidified incubator containing 5% CO_2_. HT22, J774A.1, MIN6, and MDA-MB-231 cells were grown as adherent cultures in DMEM, while Jurkat cells were grown in suspension in RPMI. The growth medium was supplemented with 100 U/mL penicillin,100 µg/mL streptomycin, and FBS (10-15%). GlutaMAX™ and sodium pyruvate were used as additional supplements in all cell lines except HT22. HT22, J774A.1, MDA-MB-231, and MIN6 cells were cultured in 4.5 g/L glucose. Jurkat cells were cultured in 1 g/L glucose. For passaging, adherent cells were detached from the plate using trypsin or a cell scraper (J774A.1 cells).

Lentivirus was produced using a combination of three plasmid vectors: a transfer plasmid encoding Aqp1 or GFP, a packaging plasmid, and an envelope plasmid encoding the VSV-G protein to confer broad cell-type tropism. To produce lentivirus, 22 µg of the transfer plasmid, 22 µg of the packaging plasmid, and 4.5 µg of the envelope plasmid were mixed and delivered to 293T cells by transient transfection with polyethyleneimine. Approximately 24 h after transfection, cells were treated with sodium butyrate (10 mM) to enhance the expression of viral genes. Viral production was allowed to continue for another 72 h before collecting the spent media and precipitating the lentiviral particles using Lenti-X™ concentrator. The concentrated lentivirus was resuspended in 200 µL PBS and stored as aliquots at -80 °C.

For lentiviral transduction, cells were grown in individual wells of a six-well plate to 70% confluence, aspirated to remove spent media, and incubated with lentiviral particles resuspended in ∼ 1 mL of media containing 8 µg/mL polybrene. Cells were spinfected by centrifuging the six-well plates at 1048 × g at 30 °C for 90 min. Following spinfection, the plates were returned to the incubator for two days. Next, the cells were transferred from to a 10-cm plate and grown to 70% confluence. Stably transduced green fluorescent cells were enriched using FACS (Sony SH800 or Sony MA900 sorter), and the enriched populations were grown back out and stored as cryo-stocks until further use.

### Magnetic resonance imaging

Cells were seeded 24-48 h prior to MRI and grown to full confluence in 10-cm tissue culture plates. In preparation for imaging, cells were harvested from the plate, centrifuged at 350 × g for 5 min, and resuspended in 200 µL of PBS. The resuspended cells were transferred to a microcentrifuge tube, centrifuged at 500 × gfor 5 min, and the supernatant was carefully aspirated. The wash step was repeated one more time before centrifuging the cells (500× g, 5 min) to form a pellet. Tubes containing cell pellets were placed in a water-filled agarose mold (1% w/v) that was housed in a custom 3D-printed MRI phantom.

All MRI experiments were performed at ambient temperature using a 7 T vertical-bore MRI scanner (Bruker) equipped with a 66 mm diameter volume coil. Stimulated echo diffusion-weighted images of cell pellets were acquired in the axial plane using the following parameters: echo time, T_E_ = 18 ms, repetition time, T_R_ = 1000 ms, gradient duration, δ = 5 ms, gradient separation, Δ = 300 ms, matrix size = 128 × 128, field of view (FOV) = 5.08 × 5.08 cm^2^, slice thickness = 1-2 mm, number of averages = 5, and four effective b-values in the range of 1-3 ms/µm^2^. The diffusion-weighted intensity at a given b-value was estimated by computing the mean intensity inside a manually drawn region of interest (ROI) encompassing the axial cross-section of a cell pellet. The slope of the logarithmic decay in the signal intensity versus the effective b-value was used to calculate the apparent diffusion coefficient. To generate diffusion maps, apparent diffusivity was computed for each voxel in the ROI. The ensuing image was smoothed using a median filter and pseudo-colored according to an 8-bit color scale. Least-squares regression fitting was performed by using the “fitnlm” function in Matlab (R2022b).

### Cell viability assays

Aqp1- and GFP-expressing cells were seeded in 96-well plates at approximately 10,000 cells per well. Cell counting was performed manually using a hemocytometer or an automated cell counter (Countess® II). Cell viability and toxicity assays were performed by measuring intracellular ATP content, MTT bioreduction, and caspase 3/7 activation, using commercially available reagents, namely CellTiter-Glo®, CellTiter 96® AQ_ueous_, CellTiter-Blue®, and Caspase-Glo® 3/7, following the manufacturer’s instructions. Absorbance and luminescence were measured using a Tecan Spark® Microplate Reader. The integration time for luminescence was set at 1 s. The caspase 3/7 activation measurements were normalized to the viable cell count based on the total cellular ATP or MTT reduction.

### Quantitative phase imaging

Quantitative phase imaging was performed using a Nanolive 3D cell explorer to determine the dry mass, sphericity, and volume of Aqp1- and GFP-expressing cells. For phase imaging, cells were seeded at a 1:24 split ratio and grown for ∼ 24 h in 35 mm glass bottom plates (Ibidi). This split ratio was chosen to ensure that the cells did not become too confluent at the time of imaging, which would hinder the accurate segmentation of individual cells. Tomographic phase images (0.202 µm^2^ lateral resolution, 0.363 µm z-resolution) were obtained from multiple regions of the plate, and cells were rendered using the “Surfaces” model in Imaris 9.0 (Oxford Instruments). Cells close to the image boundaries and those that could not be clearly differentiated from neighboring cells were excluded from the analysis. Imaris was used to compute cell volume (*V*) and surface area (*S*), which were then used to estimate sphericity (Ψ) as 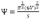. The cell mass was calculated from the refractive index tomogram using an established approach. Briefly, the refractive index difference is related to the phase shift or change in optical path length (*OPL*) as follows: 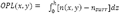 coordinate along the optical axis, *h* is the thickness of the cell, *n(x, y)* is the refractive index of the cellular material at planar coordinates *(x, y)*, and *n*_sur_ is the refractive index of the surrounding medium. The integral of the optical path difference over the plane of a segmented cell is related to dry mass (*m*) by the equation: 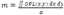 where *α* = 0.18 *µm*^3^/*pg* **is a** known constant for most eukaryotic cells ^71^.

### Unfolded protein response (UPR) assays

Aqp1- and GFP-expressing cells were cultured in six-well plates for 24 h prior to experimentation. To induce ER stress, the culture medium was supplemented with tunicamycin at a final concentration of 2.5 µg/mL for 4 h. Cells were lysed in Buffer RLT (Qiagen) supplemented with 1% β-mercaptoethanol and the lysate was centrifuged in QIAshredder columns to shear genomic DNA. Total RNA was extracted according to the manufacturer’s (Qiagen) protocol. Next, 1 µg of RNA was reverse-transcribed using iScript™ cDNA synthesis reagents according to the manufacturer’s protocol. The cDNA was diluted 10-fold in nuclease-free water and gene-specific quantitative PCR (qRT-PCR) was carried out using Power Up™ SYBR™ Green Master Mix and 20 ng of cDNA template, according to the manufacturer’s protocol. qRT-PCR was performed using a CFX96 Touch Real-Time PCR Detection System (Bio-Rad). Cycle threshold values (*C*_*t*_) were determined by regression fitting using CFX Maestro Software (Bio-Rad). Primers (**Table S2**) were designed using Primer3, ensuring that the sequences were intron-flanking, produced an amplicon between 50 and 250 bp, and had a melting temperature (*T*_*m*_) of 60 °C. The *T*_*m*_ was experimentally verified by thermal gradient PCR. To ensure that the amplification was specific to the target gene, melt curve analysis was performed, and the expected amplicon size was further confirmed using agarose gel electrophoresis. Additionally, we performed qPCR on serial dilutions of a cDNA sample (pooled across treatment conditions) to assess primer efficiency based on the slope of *C*_*t*_ vs. logarithmic dilution. Changes in expression of key UPR-associated genes were quantified using the 2^−*ΔΔC*^_*t*_ method^72^ with GAPDH and actin serving as housekeeping genes for mouse and human cell lines respectively.

### Phagocytosis assay

Supported lipid bilayer-coated silica beads were prepared as described before^73^. Briefly, chloroform-dissolved lipids were mixed in the following molar ratios: 96.8% POPC (Avanti, Catalog # 850457), 2.5% Biotinyl Cap PE (Avanti, Catalog # 870273), 0.5% PEG5000 PE (Avanti, Catalog # 880230), and 0.2% atto390-DOPE (ATTO-TEC GmbH, Catalog # AD 390–161), and dried under argon gas overnight to remove the chloroform. The dried lipids were resuspended to 10 mM in PBS (pH 7.2) and stored under argon gas. To form small unilamellar vesicles (SUVs), the lipid mixture was subjected to 30 freeze-thaw cycles and stored at -80 °C under argon gas. Immediately before use, the SUVs were filtered through 0.22 µm PTFE syringe filters to remove lipid aggregates. To form supported lipid bilayers, 8.6 × 10^8^ silica beads (10 µL of 10% solids, 4.89 µm mean diameter, Bangs Laboratories) were washed twice with water and twice with PBS by centrifugation at 300 × g and decanting. The beads were then mixed with 1 mM SUVs in PBS, vortexed for 10 s at medium speed, covered with foil, and incubated in an end-over-end rotator at room temperature for 0.5-2 h to allow lipid bilayers to form on the beads. The beads were washed three times with PBS to remove excess SUVs and resuspended in 100 µL of 0.2 % casein in PBS for 15 min at room temperature to block nonspecific binding. Anti-biotin monoclonal mouse IgG1κ (Jackson Immuno Labs, Cat#200-602-211) was added to the beads at concentrations ranging from 0 – 10 nM and incubated for 15 min at room temperature with end-over-end mixing. The beads were washed three times to remove unbound IgG and resuspended in 100 µL PBS containing 0.2 % casein.

For the bead engulfment assay, ∼ 50,000 Aqp1-or GFP-expressing J774 cells were plated in individual wells of a 96-well glass bottom plate (MatriPlate™) approximately 12-24 h prior to the experiment. Next, the cells were washed four times with engulfment imaging media (20 mM HEPEs, 135 mM NaCI, 4 mM KCl, 10 mM glucose, 1 mM CaCl_2_, 0.5 mM MgCl_2_), leaving ∼ 100 µL media between washes, and finally leaving the cells in 300 µL media. Approximately 8 ×10^5^ beads were added to each well and engulfment was allowed to proceed for 45 min in a humidified incubator (37 °C, 5 % CO_2_). A minimum of 36 Images were collected using a spinning disk confocal microscope (Nikon Ti2-E inverted microscope with a Yokagawa CSU-W1 spinning disk unit and an ORCA-Fusion BT scientific CMOS camera) with a 40X 0.95 NA Plan Apo air objective. The microscope was operated using the NIS-Elements software platform (Nikon). Images were analyzed using CellProfiler^74^. All images were cropped to remove 150 pixels around the edges and background was subtracted. Macrophage cell bodies were identified from the GFP channel using a global minimum cross-entropy threshold. The phagocytic index was estimated from average integrated fluorescence intensity of atto390-labeled lipids per macrophage. Results were validated by a blinded analyzer manually counting the number of beads in 100 macrophages per condition.

### Insulin secretion assay

Aqp1- and GFP-expressing MIN6 cells were seeded in individual wells of a 24-well plate at approximately 120,000 cells per well (manually counted using a hemocytometer). Four days after seeding, the culture media was aspirated and cells were washed twice with low-glucose Krebbs-Ringer bicarbonate/HEPES medium (KRBH) comprising 135 mM NaCl, 3.6 mM KCl, 2 mM NaHCO_3_, 0.5 mM NaH_2_PO4, 0.4 mM MgCl_2_, 1.5 mM CaCl_2_, 10 mM HEPES, 2 mM glucose, and 1% BSA (pH 7.4). The cells were subsequently incubated at 37 °C for 2 h in low-glucose KRBH medium, washed twice as before, and incubated at 37 °C in KRBH media containing either 2 mM (low) or 20 mM (high) glucose. One hour later, 450 µL of supernatant was collected from each well, centrifuged at 1500 × g for 5 min at 4 °C, and stored at -80 °C. Next, the cells were lysed using RIPA Lysis Buffer, and the lysates were stored at -80 °C. The insulin concentration in the thawed supernatant was determined using a mouse insulin ELISA Kit following manufacturer’s recommendations to obtain a dynamic range of 0.1 – 12.8 ng/mL insulin. A standard curve was generated by measuring known quantities of purified insulin and fitting the curve to a quadratic model. Insulin concentrations were normalized to total cellular protein, which was measured in thawed lysates using the bicinchoninic acid (BCA) assay.

### Flow cytometry assay for immune stimulation

We first confirmed the expression of various immune receptors, including CD3, CD4, and CD45 in native Jurkat cells by staining with the following antibodies, namely anti-human CD3-PE (UCHT1, 0.024 µg), anti-human CD4-PE-Cy7 (OKT4, 0.024 µg), and anti-human CD45 eFluor® 450 (HI30, 0.2 µg),. Staining was performed by pelleting Jurkat cells at 500 × g, washing with staining buffer (PBS containing 1% FBS)and incubating with the respective antibodies for 20 min (200 µL staining buffer, 2 µL respective antibody). To avoid cross-staining, each receptor was independently stained. Next, Aqp1- and GFP-expressing Jurkat cells (∼500,000 cells/mL) were stimulated using anti-human CD3/CD28 magnetic Dynabeads® at a 4:1 bead-to-cell ratio (2 uL). After 24 h of stimulation, Jurkat cells were collected and de-beaded using a magnetic rack for 1-3 minutes. The de-beaded cells were pelleted at 500 × g by centrifugation, washed with PBS containing 1% FBS, and stained with mouse anti-human CD3-PE antibody for 20 min. After CD3 staining, the cells were washed as before and stained with anti-mouse CD25 eFluor® 450 (PC61.5, 0.1 µg). Stimulation was assessed by using flow cytometry (Attune NxT Flow Cytometer) to quantify the percentage of CD3+ cells that were also CD25+. Flow cytometry data were analyzed using FlowJo. The gating strategy is illustrated in **Figure 4d** and **Supplementary Fig. 3**. Percentage of CD25+ cells was calculated relative to the total CD3+ population as *stimulation*%= *CCCC*3^+^*CCCC*25^+^/[*CCCC*3^+^*CCCC*25^+^ +*CCCC*3^+^*CCCC*25^−^].

### Matrigel invasion assay

MDA-MB-231 cells were grown to 70% confluence and ∼ 50,000 cells were centrifuged at 350 × g for 5 min, decanted, and resuspended in 10 µL Matrigel containing Phenol Red. The resuspended cells were deposited in the form of single drops in individual wells of a six-well plate and incubated at 37 °C for 20 min to allow the Matrigel to solidify. Next, 2 mL of medium was added to each well, and the plate was returned to the incubator. The cells were imaged every 24 h using a scanning confocal microscope (Leica SP8). Both transmitted light and fluorescent images were acquired using a 10 × 10 grid of 1024 × 1024 pixel tiles at 10X magnification to capture the Matrigel drop and the surrounding medium. Fluorescence imaging was performed using 488 nm light for excitation and emission was measured between 498 and 592 nm. Imaging was continued for up to six days, beyond which the Matrigel drops began to detach from the plate. For image analysis, Fiji was first used to manually draw boundaries around the Matrigel border and zero-fill the pixels located inside the ROI, thereby eliminating cells that did not migrate out of the droplet. Next, ilastic was used to automatically segment cells located outside the Matrigel border, using images obtained at intermediate times (e.g., day 4) as the training set. Finally, the percentage of all pixels in the image that corresponded to escaped cells was computed using Fiji and converted to area units by multiplying by the total imaged area (113.5 mm^2^).

### Statistical analysis

Experimental data are summarized by their mean and standard error of mean obtained from multiple (*n* ≥ 3) biological replicates. All tests are 2-sided with the exception of the tunicamycin-based UPR assay (Fig. 3b), which was 1-tailed given the directionality of tunicamycin’s effect on UPR was known a priori. The qRT-PCR data were analyzed by performing a Student’s t-test onthe *ΔΔC*_*t*_ values. A *P-*value of less than 0.05 taken to indicate statistical significance.

## Ethics approval and consent to participate

Not applicable

### Consent for publication

Not applicable

## Availability of data and materials

The datasets used and/or analysed during the current study are available from the corresponding author on reasonable request.

## Authors’ Contributions

**Austin D.C. Miller**: Conceptualization, Methodology, Investigation, Validation, Formal analysis, Writing - Original Draft, Writing - Review & Editing. **Soham Chowdhury, Hannah Ghasemi, Wyatt Miller**: Methodology, Investigation, Validation, Formal analysis, Writing - Original Draft, Writing - Review & Editing.: **Hadley Hanson, Sarah K. Linderman**: Investigation, Validation. **Meghan Morrissey, Chris Richardson, Brooke Gardner**: Methodology, Resources, Writing- Review & Editing, Funding acquisition. **Arnab Mukherjee:** Conceptualization, Methodology, Formal analysis, Resources, Data Curation, Writing - Original Draft, Writing - Review & Editing, Visualization, Supervision, Project administration, Funding acquisition.

## Acknowledgements

We thank Dr. Harun F. Ozbakir for helpful discussions and Dr. Ben Lopez for providing excellent technical support in the microscopy experiments.

