## Supplementary Information for "Engineering water exchange is a safe and effective method for magnetic resonance imaging in diverse cell types"

### Supporting Figures

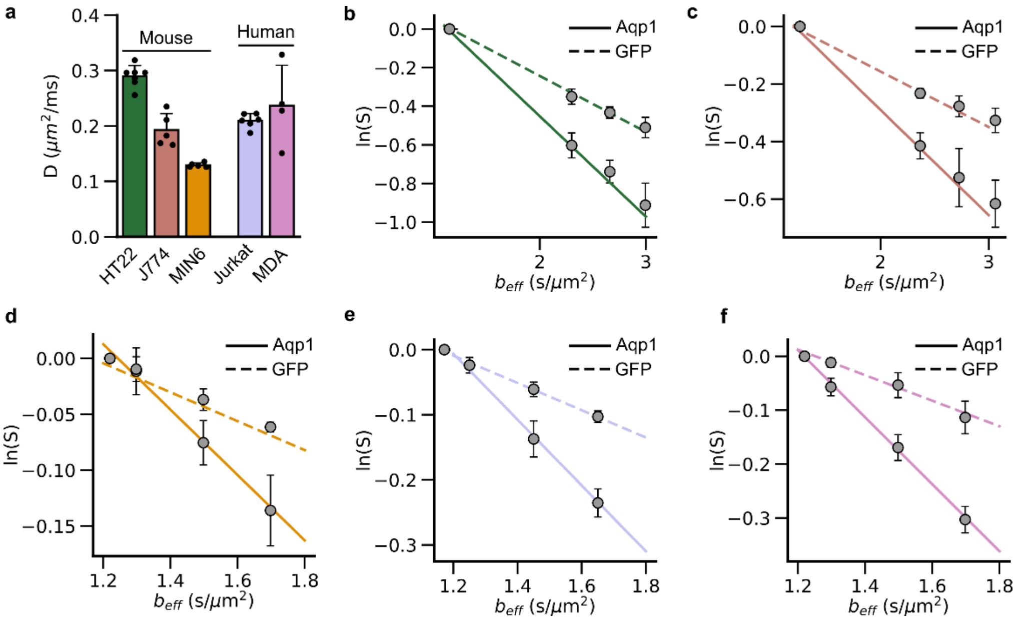


**Figure S1: Measurement of diffusivities in wild-type and Aqp1-expressing cells**. **a**, Baseline diffusivities were measured in cells transduced to express GFP. Aqp1 expression increases diffusivity over baseline, leading to a faster decay in diffusion-weighted signal intensity ($S$) with effective b-value ($b_{eff}$) in all five cell types: **b**, HT22, **c**, J774, **d**, MIN6, **e**, Jurkat, and **f**, MDA-MB-231 cells. The solid and dotted lines represent the first-order decay in signal intensity in Aqp1- and GFP-expressing cells, respectively. Error bars represent s.e.m. ($n \geq 4$ biological replicates). All MRI data were acquired at 7 T, using a diffusion time of 300 ms.

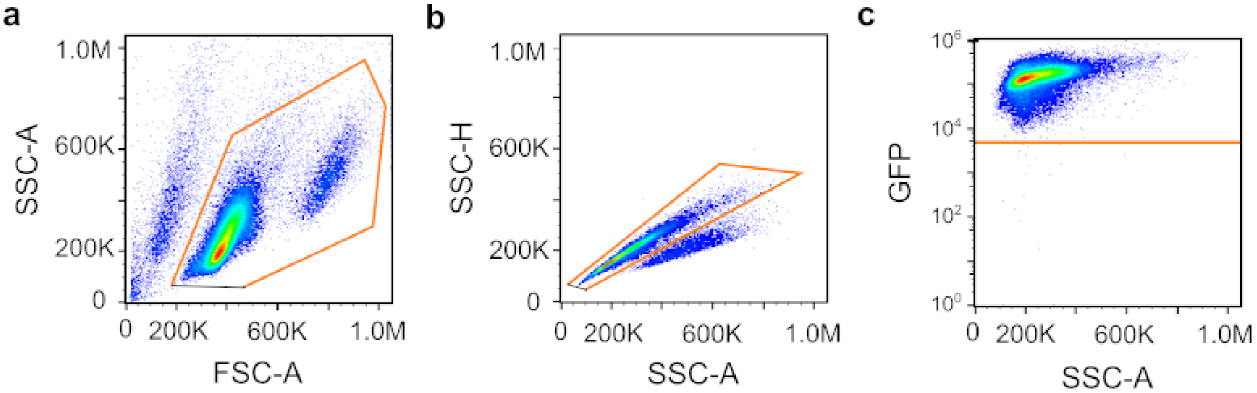


**Figure S2: Gating strategy to select reporter-expressing following lentiviral transduction**. Representative flow cytometry analysis depicting the gating strategy used to generate uniform populations of reporter-expressing cells by gating for a, viability b, singlet cells, and c, GFP expression. FSC-A indicates forward scatter area. SSC-H and SSC-A respectively denote side scatter height and side scatter area.

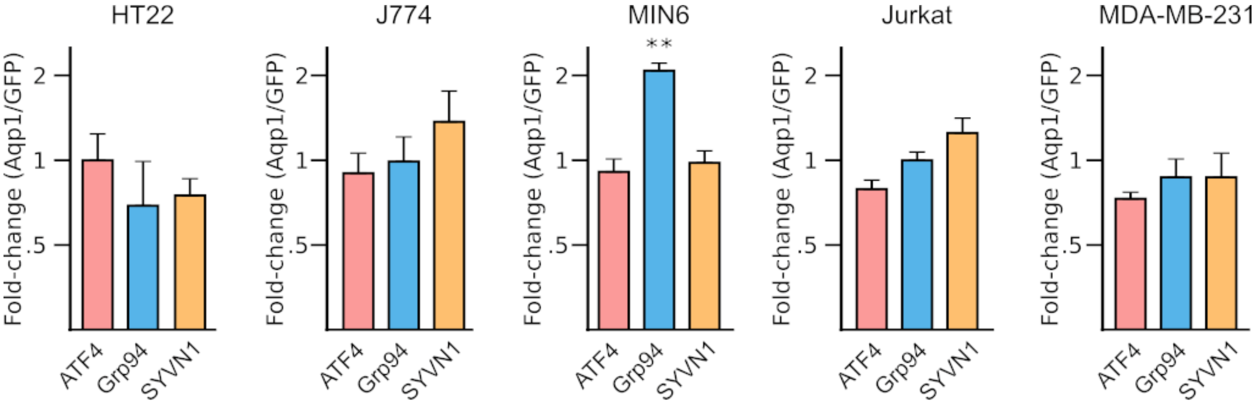


**Figure S3: Effect of Aqp1 expression on the unfolded protein response**. Fold changes in the expression of UPR-associated genes, including ATF4, Grp94, and SYVN1 in Aqp1-expressing cells relative to GFP controls. Error bars represent the s.e.m. from *n* = 3 biological replicates. ** *P*-value < 0.01 (2-sided t-test). GAPDH and actin were used as housekeeping genes for the mouse and human cell lines, respectively.

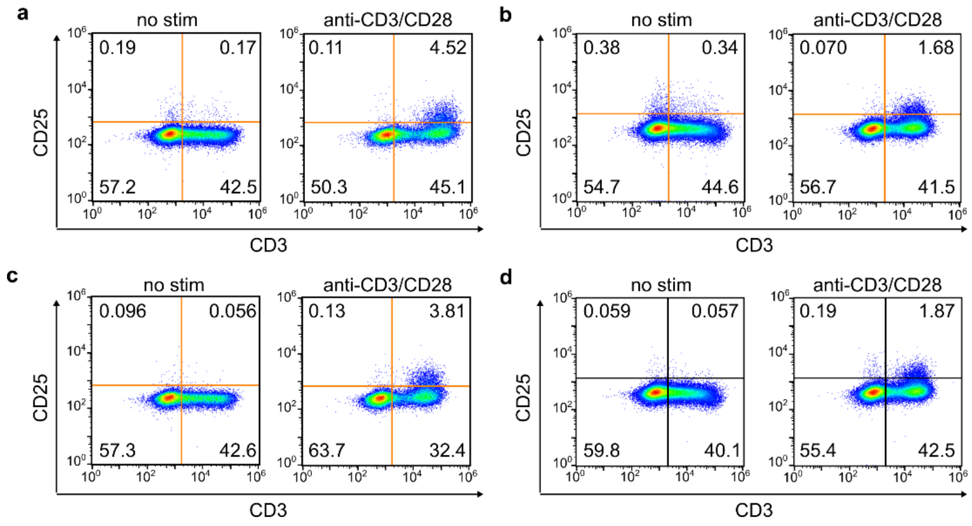


**Figure S4. Flow cytometry analysis of CD25 and CD3 surface expression in Jurkat cells.** To stimulate CD25 expression, Jurkat cells were incubated with anti-human CD3/CD28 beads at a 5:1 bead-to-cell ratio for 24 h. (**a,c**) Aqp1- and (**b,d**) GFP-expressing Jurkat cells were analyzed for surface expression of CD3 (pan T-cell marker) and CD25 (activation marker) using flow cytometry. The numbers in each quadrant denote the fraction of cells showing surface expression of CD3, CD25, both CD3 and CD25, or neither marker.

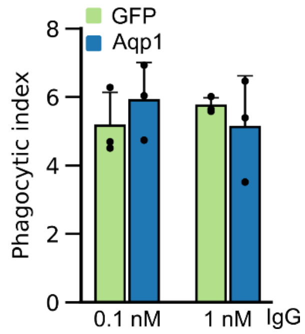


**Figure S5: Phagocytic activity of Aqp1- and GFP-expressing macrophage cells**. Phagocytic index was computed following exposure of J774 cells to lipid-coated silica beads with 0.1 or 1 nM IgG1κ, for 45 min.

**Table S1**: General characteristics of cell lines used in the study

| **Cell line** | **Type** | **Species** | **Sex** | **Culturing** |
|---|---|---|---|---|
| HT22 | hippocampal neuron | mouse | unknown | adherent |
| J774A.1 | macrophage, monocyte | mouse | female | adherent |
| MIN6 | insulinoma, pancreatic beta cell | mouse | unknown | adherent |
| Jurkat E6-1 | T lymphoblast | human | male | suspension |
| MDA-MB-231 | epithelial, triple negative breast cancer | human | female | adherent |

**Table S2**: Oligonucleotide primers for qRT-PCR

| Gene | Sequence |
|---|---|
| MmXBP1s_F | TGAGTCCGCAGCAGGTG |
| MmXBP1s_R | TCCTTCTGGGTAGACCTCTGG |
| MmATF4_F | GAAACCTCATGGGTTCTCCA |
| MmATF4_R | AGAGCTCATCTGGCATGGTT |
| MmCHOP_F | CACCACACCTGAAAGCAGAA |
| MmCHOP_R | GGACGCAGGGTCAAGAGTAG |
| MmBip_F | GGACAAGAAGGAGGATGTGG |
| MmBip_R | TGATCGTTGGCTATGATCTCC |
| MmSYVN1_F | AACCCCACTGAAGAGACTGC |
| MmSYVN1_R | CTCGGGAAGCTCCTCTACAA |
| MmGrp94_F | GTCGGGAAGCAACAGAGAAG |
| MmGrp94_R | TGCCAGACCATCCATACTGA |
| MmGAPDH_F | CCACCCAGAAGACTGTGGAT |
| MmGAPDH_R | CACATTGGGGGTAGGAACAC |
| HsXBP1s_F | GGAGTTAAGACAGCGCTTGGG |
| HsXBP1s_R | CTGCACCTGCTGCGGAC |
| HsATF4_F | TCAAACCTCATGGGTTCTCC |
| HsATF4_R | GTGTCATCCAACGTGGTCAG |
| HsCHOP_F | AGCGACAGAGCCAAAATCA |
| HsCHOP_R | CAGTGTCCCGAAGGAGAAAG |
| HsBip_F | CATGGTTCTCACTAAAATGAAAG |
| HsBip_R | GCTGGTACAGTAACAACTG |
| HsSYVN1_F | CCAACATCTCCTGGCTCTTT |
| HsSYVN1_R | GTCAGGATGCTGTGATAGGC |
| HsGrp94_F | GTCGGAAAAGTTTGCCTTCC |
| HsGrp94_R | TGACATGCAGCAGGTTCTTC |

| | |
|---|---|
| HsACTB_f | TTGGCAATGAGCGGTTCC |
| HsACTB_r | GTTGAAGGTAGTTTCGTGGATG |
